# Dispersal predicts hybrid zone widths across animal diversity: Implications for species borders under incomplete reproductive isolation

**DOI:** 10.1101/472506

**Authors:** Jay P. McEntee, J. Gordon Burleigh, Sonal Singhal

## Abstract

Hybrid zones occur as range boundaries for many animal taxa. One model for how hybrid zones form and are stabilized is the tension zone model. This model predicts that hybrid zones widths are determined by a a balance between random dispersal into hybrid zones and selection against hybrids, and it does not formally account for local ecological gradients. Given the model’s simplicity, it provides a useful starting point for examining variation in hybrid zone widths across animals. Here we examine whether random dispersal and a proxy for selection against hybrids (mtDNA distance) can explain variation in hybrid zone widths across 135 hybridizing taxon pairs. We show that dispersal explains >30% of hybrid zone width variation across animal diversity and that mtDNA distance explains little variation. Clade-specific analyses revealed idiosyncratic patterns. Dispersal and mtDNA distance predict hybrid zone widths especially well in reptiles, while hybrid zone width scaled positively with mtDNA distance in birds, opposite predictions. Lastly, the data suggest that lower bounds on hybrid zone widths may be set by dispersal and the extent of molecular divergence, suggesting that hybrid zones are unlikely to form in restricted geographic spaces in highly dispersive and/or recently diverged taxa. Overall, our analyses reinforce the fundamental importance of dispersal in hybrid zone formation, and more generally in the ecology of range boundaries.

## Introduction

The edge of a taxon’s geographic range often abuts the edge of the range of another, closely-related taxon (Case and Taper 2000). If reproductive isolation between these taxa is incomplete, hybrid zones can form. Here, we adopt Harrison’s definition of hybrid zones as “interactions between genetically distinct groups of individuals resulting in at least some offspring of mixed ancestry” (Harrison 1990), with genetically pure populations found outside of the zone. The proportion of hybrid offspring in hybrid zone populations varies from very low (1.3% in *Triturus* newts; (Arntzen and Wallis 1991)) to very high (93.3% between phylogeographic lineages in the Australian lizard *Lampropholis coggeri*; (Singhal and Moritz 2013)). Hybrid zones most often form when previously-isolated populations come into secondary contact due to changing range boundaries (Remington 1968) but may also form in place while populations diverge with gene flow (Haldane 1948; Endler 1977; Nosil 2012). Often, these regions of contact are narrow relative to the distributions of pure populations, even when they extend along significant swaths of a species’s range (Figs. 1 and 2).

**Figure 1.**
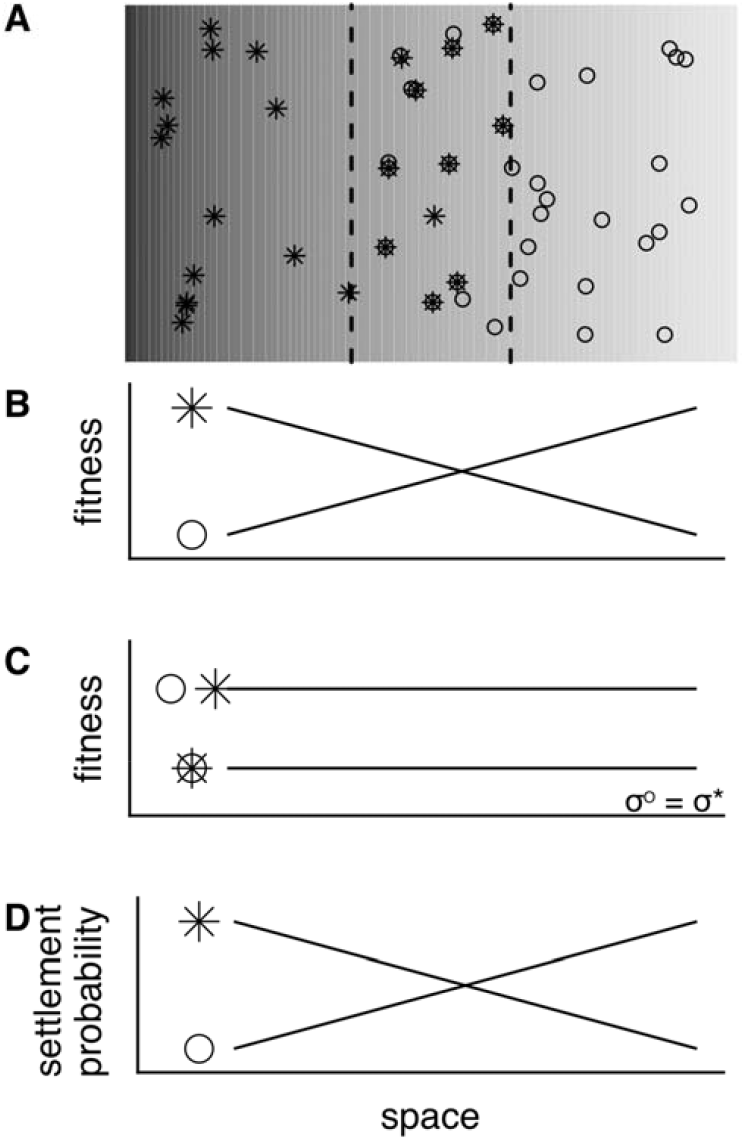
(A) Depiction of a clinal hybrid zone, where two differentiated taxa (circles and crosses) meet and form hybrids (crosses inside circles, here generally indicating individuals of mixed ancestry). The dashed lines indicate the approximate boundaries of where hybrids are formed. Models to explain the stabilization of these hybrid zones are not mutually exclusive, but invoke different processes. In (B), the fitness of the two taxa vary inversely along an ecological gradient, with hybrids formed in the area where they have similar fitness. The environmental gradient is indicated by the shading in (A). (C) depicts the tension zone model, in which fitness is independent of position along a transect across the hybrid zone, but where hybrids always have lower fitness than ‘pure’ individuals. Stabilization occurs as a balance between selection against hybrids and dispersal rate σ, which is generally modeled as being similar between taxa. (D) depicts a habitat choice model where taxa prefer to live or breed in particular environments, and hybrids are formed where the probability of breeding is similar.

**Figure 2.**
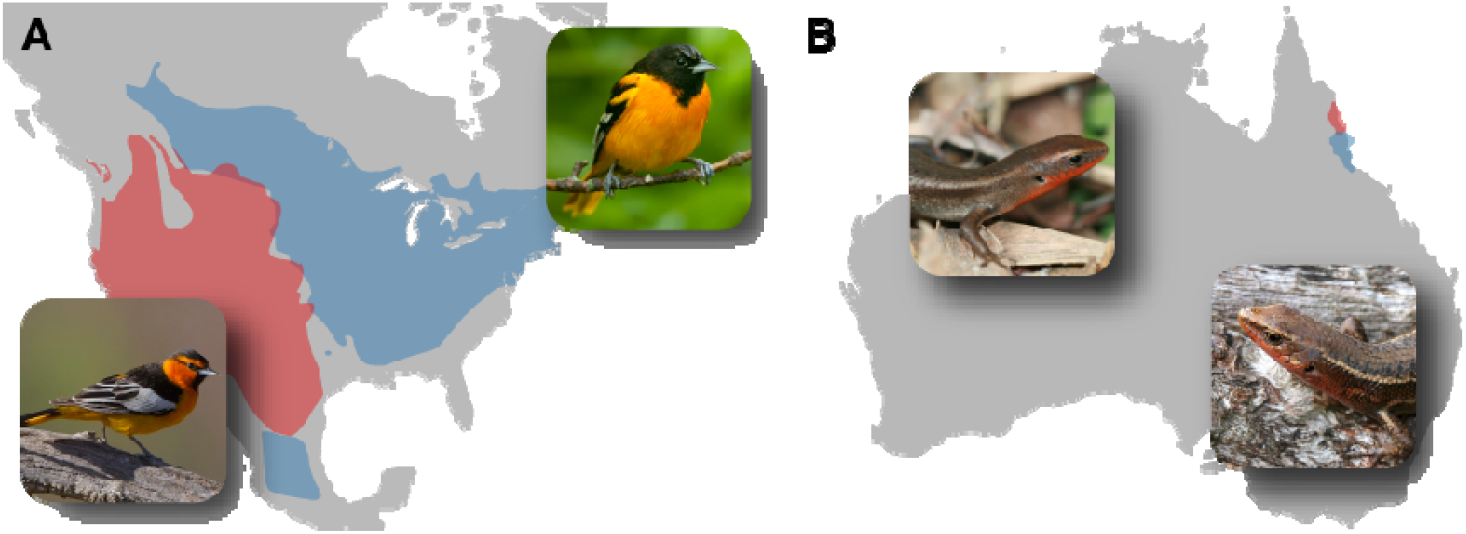
Geographic ranges and species photos for two of the hybrid zones included in this study, A. *Icterus bullocki* (red, L) and *I. galbula* in North America (blue, R) and B. *Carlia crypta* (red, L) and *C. rubrigularis* (blue, R) in Australia. The *I. bullocki* / *I. galbula* hybrid zone is geographically extensive and occurs between temperate, high-dispersing, morphologically differentiated species. In contrast, the *C. crypta* / *C. rubrigularis* hybrid zone is geographically narrow and occurs between tropical, low-dispersing, morphologically cryptic species. These two hybrid zones exemplify some of the diversity in hybrid zones. Image credits: *I. bullocki* (photograph by Gregory Smith; distributed under a CC-BY 2.0 license), *I. galbula* (photograph by Laura Gooch; distributed under a CC-BY 2.0 license), *C. crypta* (photograph by Ben Phillips), *C. rubrigularis* (photograph by Sonal Singhal).

Hybrid zones can form across a spectrum of divergence stages between taxa. At one extreme, hybrid zones can occur between taxa that vary at only a few genes throughout the genome, such as in golden- and blue-winged warblers (Toews et al. 2016). At another, hybrid zones form between species that are millions of years diverged and exhibit marked genomic and phenotypic divergence, such as the common and bay mussel (Stuckas et al. 2009). In fact, hybrid zones regularly occur between recognized taxa that are considered “good species”, even though they regularly interbreed. Hybridizing taxa have been diagnosed widely across the tree of life and geographic regions ((Barton and Hewitt 1985), Fig. S1), and studies of these diverse hybrid zones have been valuable for understanding how species form and are maintained. For example, these studies have shown that increased molecular divergence between taxa correlates with increased selection against hybrids (Singhal and Moritz 2013), that genes putatively under direct selection in hybrids show reduced introgression (Rieseberg et al. 1999), and that selection against hybrids can drive the evolution of pre-mating isolation (Hoskin et al. 2005).

While hybrid zones have long been the subject of evolutionary inquiry (Endler 1977; Barton and Hewitt 1985; Harrison 1990), hybrid zones remain underappreciated in ecology, particularly in the study of geographic range limits. Geographic range limits are thought to arise from a number of factors, including barriers to dispersal, abiotic limits, and biotic interactions (Case and Taper 2000; Case et al. 2005; Hochkirch et al. 2007; Sexton et al. 2009; Jankowski et al. 2013; Weber and Strauss 2016). These biotic interactions can include hybridization ((Case and Taper 2000; Case et al. 2005; Hochkirch et al. 2007; Sexton et al. 2009; Jankowski et al. 2013; Weber and Strauss 2016)), but most ecological work on range boundaries has focused on other factors. However, given the ever-increasing evidence for the prevalence of hybridization across the animal tree of life (Mallet et al. 2016), hybridization is likely to be important in the formation of many range boundaries even where hybrids have yet to be found (Levin 2006). Thus, the extensive existing body of empirical and theoretical work on hybrid zones, though largely focused on evolutionary questions, may also yield important ecological lessons.

For a number of reasons, hybrid zones are powerful systems through which to study range limits. First, because hybrid zone studies typically sample densely at species’ edges, they provide high-resolution data for the analysis of range boundaries. Second, hybrid zone data are typically analyzed across a fairly small set of standardized approaches. Because these approaches are somewhat standardized, we can compare the spatial scale of range limits across phylogenetic distant relatives, like mammals and insects. Third, efforts to re-survey hybrid zones or make estimates across different sampling periods from museum collections have generated fine-scale estimates of hybrid zone change. This work may allow us to better understand how range limits vary over time, and what factors drive changes in hybrid zone width and location (Dasmahapatra et al. 2002; Buggs 2007; Taylor et al. 2015).

Much of our understanding of how hybridization can set range limits derives from explorations of theoretical models of hybrid zone stabilization. With respect to ecology, these models can be divided into two classes: those that require ecological gradients or habitat transitions (Endler 1977; Goldberg and Lande 2007; Armsworth and Roughgarden 2008; Price and Kirkpatrick 2009) and those that do not (Bazykin 1969; Barton and Hewitt 1985; Case et al. 2005; Goldberg and Lande 2007). Empirically, hybrid zones are found centered along environmental gradients (Endler 1977), such as changes in soil composition (Patton 1993), and also in the apparent absence of such gradients (Barton and Hewitt 1985). In hybrid zone models requiring ecological variation across space, each parental taxon is adapted to some environment, and shows decreased fitness elsewhere (Figure 1). An example is mosaic hybrid zones (Harrison 1990), in which taxa map precisely onto patches of interdigitated habitats where they come into contact (Ross and Harrison 2002). In hybrid zone models associated with ecological variation, hybrid offspring are typically limited to the ecotone, where their fitness is similar to that of parental taxa (Fig. 1). In some instances, hybrid fitness may even be higher in intermediate (or “hybridized”) environments (Moore 1977). In contrast, in models without ecological gradients, fitness does not depend on local environmental conditions (Bazykin 1969; Barton and Hewitt 1985). One such model is the tension zone model (Key 1968; Bazykin 1969; Barton and Hewitt 1985), in which hybrids show reduced fitness compared to their parents, either because of intrinsic selection (e.g. Dobzhansky-Muller incompatibilities resulting in hybrid breakdown; (Dobzhansky 1936; Muller 1942), or because hybrid traits have low fitness in all environments (Mallet et al. 1998). Importantly, the location of a tension zone can be random with respect to geographic space and local ecological gradients, though it is predicted to move towards regions of reduced dispersal (Barton and Hewitt 1985; Goldberg and Lande 2007). Of course, both ecological variation and selection against hybrids independent of local ecology can be important within a single hybrid zone (Bronson et al. 2003; Taylor et al. 2014).

Ideally, to understand how range boundaries associated with hybridization are typically formed and maintained, we could reconstruct the conditions under which these boundaries are formed. Through this, we could understand how ecological transitions, taxon fitness across such transitions, and taxon habitat preference -- or the lack thereof -- interact to structure hybrid zones and enforce range boundaries of hybridizing taxa (Fig. 1). Optimally, we would experimentally investigate the fitness of taxa and their hybrids in transects across hybrid zones (Mallet and Barton 1989; Prada and Hellberg 2013). In most animals, these data do not exist, and would be extremely time- and cost-expensive - if not implausible - to collect. Absent these more direct approaches, we can pursue an alternate path to investigate our questions: we can examine whether variation in hybrid zone widths can be predicted by the factors that stabilize hybrid zones in theoretical models. To make our predictions, we focus on the tension zone model both because it has been extensively applied in the hybrid zone literature (Barton and Gale 1993; Gay et al. 2008) and because the model assumes a homogeneous environment, making it a useful null model for how hybrid zones might form and stabilize with respect to local ecology. At its simplest form, the tension zone model predicts that hybrid zones are stabilized by just two forces, random dispersal and selection against hybrids. The balance between these two forces determines the width of the hybrid zone boundary, which is expected to be stable through time under a number of demographic and ecological scenarios (Case et al. 2005; Goldberg and Lande 2007).

In this study, we address a central question in the ecology of hybrid zones: which factors determine the widths of hybrid zones? To answer this question, we conduct a meta-analysis of 135 pairs of hybridizing animal taxa. In particular, we explore how well hybrid zones -- and thus range limits -- can be explained by the tension zone model by testing if dispersal and selection against hybrid taxa explains hybrid zone widths. More generally, this study represents the first quantitative meta-analysis of hybrid zones since (Barton and Hewitt 1985) and thus provides a novel summary of patterns from hybrid zones in the DNA sequencing era.

## Methods

### Literature Review

To identify animal hybrid zones for inclusion in this meta-analysis, we surveyed the literature using Google Scholar and Web of Knowledge on two separate occasions: 6 - 8 February 2016 and 1 - 4 May 2017. We reviewed matches to the search term ‘hybrid zon*’ and additionally reviewed all papers that cited the four major software programs used to estimate clines (HZAR, (Derryberry et al. 2014); `Cfit7`, (Gay et al. 2008); `Analyse 1.3`, (Barton and Baird 1995); and `ClineFit`, (Porter et al. 1997)). In order to be included, we required that the studies measured geographic clines using a clearly-outlined method. These methods include formal statistical approaches like the maximum likelihood approach implemented in `Analyse`, and simpler estimates, e.g. using the estimated distance between populations with 20% and 80% allele frequency for biallelic loci. From all appropriate papers, we extracted all available data about estimated clines. We categorized whether the cline was morphological, behavioral, genetic, whether it characterized frequency differences (as for biallelic clines) or clinal changes in continuous values (as for morphometric measurements), and the width and center of the cline. Not all characters for which clines were estimated were diagnostic for the two taxa. Table S1 summarizes the full metadata recorded.

Additionally, we summarized metadata for each hybridizing pair. Most relevant to the current study, we 1) found estimates of dispersal in the literature, 2) measured a morphological proxy for dispersal for bird taxa only, and 3) either found or estimated the genetic distance of mtDNA sequences between hybridizing taxa. Details on how we collected these data appear below. Further, several hybridizing pairs were studied across multiple temporal or geographic transects. For these pairs, we also recorded data by transect. Table S2 summarizes the full metadata recorded for hybrid zones.

### Dispersal estimates

Under the tension zone model, hybrid zone clines are stabilized as a balance between the dispersal of hybridizing taxa into the hybrid zone and selection against hybrids. Accordingly, we identified a dispersal estimate for either or both taxa of the hybridizing pair or their close relative. Many of the included studies either generated their own estimates of dispersal or cited relevant estimates. For those studies that did not reference dispersal estimates, we used both Web of Knowledge and Google Scholar to identify other relevant literature that reported measures from either these species or their close relatives (i.e., congenerics). Dispersal was estimated across a number of population genetics-based and field-based methods. These methods include: using isolation-by-distance relationships to estimate the population-genetic dispersal parameter σ (Rousset 1997), using estimates of linkage-disequilibrium in the hybrid zone to estimate sigma (Barton and Gale 1993), measuring natal dispersal via direct observation of birth and breeding sites, and using mark-recapture studies to estimate movement rates.

Different methods for measuring dispersal make different assumptions. For example, population genetic dispersal estimates tend to be greater than direct field-based dispersal estimates because the latter tend to fail to adequately sample long-distance dispersal events (Koenig et al. 1996). Thus, while population genetic dispersal estimates are estimates of effective dispersal (i.e. they are a measure of how far individuals who have successfully bred have dispersed) instead of dispersal itself, they are likely a better index of dispersal itself because they better capture the effect of longer-dispersing individuals. Additionally, field-based estimates typically report dispersal as per year estimates, whereas genetic estimates are typically per generation, or the square-root of generations. Methodological differences in dispersal estimates across studies introduce error. However, dispersal in our dataset varied across >5 orders of magnitude (from .007 to 150 km; Fig. S2), and the errors introduced by methodological differences are unlikely to reach one order of magnitude. Thus, given the analytical approach we use, we do not expect these errors to result in qualitative differences in our results.

As literature-based dispersal estimates are not standardized across studies, we pursued an alternate, standardized proxy for a substantial sample of hybrid zone studies. For bird hybrid zones, we measured a morphological proxy for dispersal called the hand-wing index (HWI, (Claramunt et al. 2012)) on museum specimens. We measured HWI on bird study skins in the Florida Museum of Natural History (FMNH), the Museum of Vertebrate Zoology (MVZ) at the University of California, Berkeley, and the Natural History Museum, London (NHM). For each taxon in each hybridizing pair, we measured up to three adult individuals of each sex, dependent on the availability of specimens. Adult age was determined by plumage. To calculate a single HWI estimate per hybridizing pair, we first averaged HWI within-sex, then by taxon, and then by hybridizing pair. This approach ensured both sexes and taxa were evenly weighted in the final HWI estimate. The mean numbers of measured specimens per taxon and per hybridizing pair were 4.4±1.9SD and 6.5±3.8, respectively.

### Genetic distance estimates

As tension zones may be stabilized by selection against hybrids, we estimated mtDNA genetic distance between hybridizing taxa as a proxy for selection against hybrids. mtDNA distance generally correlates positively with divergence time (Moritz et al. 1987; Hung et al. 2016) which tends to result in stronger selection against hybrids (Coyne and Orr 1989; Sasa et al. 1998; Pereira and Wake 2009; Singhal and Moritz 2013). However, the extent of mtDNA and nuclear genetic divergence can be decoupled due to the unique biology of the mtDNA genome ((Galtier et al. 2009; Pereira et al. 2011)). In such cases, mtDNA divergence is less likely to predict selection against hybrids (Galtier et al. 2009; Pereira et al. 2011).

To estimate genetic distance between taxa, we searched GenBank for all available mtDNA sequence data for hybridizing taxa. Where sequences from different mtDNA loci were available for different sets of individuals, we sought to maximize the number of individuals sampled. We aligned sequence data using MUSCLE (Edgar 2004). We then calculated the average mtDNA distance between hybridizing taxa under the Tamura-Nei model of molecular evolution (Tamura and Nei 1993), using the `dist.dna` function in the R package `ape` (Paradis et al. 2004). If sequences for multiple loci were available for the same set of individuals, we averaged our distance estimates by length across loci. Most hybridizing pairs showed reciprocal monophyly in neighbor-joining phylogenies inferred with the mtDNA data, which suggests that most of our distance estimates were not reduced by recent introgression of mtDNA.

For nine of our taxon pairs, we were either unable to find mtDNA genetic data on GenBank, or we were unable to match the available genetic data to the hybridizing pair. For these studies, we instead include estimates of mtDNA genetic divergence as reported in the literature. For bird-only analyses (see below), we used an alternate proxy for selection against hybrids, which was the time to the most recent common ancestor (MRCA) in a large, supermatrix, species-level phylogeny, inferred using both mtDNA and nuclear loci (Burleigh et al. 2015). This proxy was only available for hybridizing bird taxa that are delineated as different species (Clements et al. 2011).

### General modeling approach

The goal of our analyses is to investigate whether dispersal and selection against hybrids can explain global variation in hybrid zone width among different hybrid zones. Hybrid zone width measures the transition between parental genotypic or phenotypic traits in the hybrid zone. We present analyses from two different general approaches to the data. In the first “all widths” approach, we analyze all reported cline estimates for each hybrid zone. This approach models the variance across widths of independent traits across hybrid zones, and the variance in widths across hybrid zones. In the second “mean widths” approach, we took the geometric mean cline width per hybrid zone as a point estimate of the hybrid zone width. In interpreting the evidence, we largely assume that hybrid zones are at dynamic equilibrium. In non-equilibrium hybrid zones, time since formation can additionally lead to variation in widths (Endler 1977).

### The “all widths” approach

The “all widths” data set included all hybrid zone cline width estimates that we could find in the literature, including where clines had been measured for multiple traits for the same hybridizing pair, either for the same transect or for different transects. Hybrid zone studies often characterize multiple clines across different molecular markers and/or phenotypic characters for the same hybrid zone transect. Variation in width among these clines is often of primary interest in evolutionary studies of hybrid zones, as variation in introgression can reveal variation in selection and recombination across traits (Anderson 1953; Bazykin 1969; Singhal and Bi 2017; Schumer et al. 2018). Width variation can also result from methodological issues, like using non-diagnostic markers to infer cline width. Here, we are less interested in variance within a hybrid zone and more focused on explaining variance among hybrid zones. Thus, to accommodate the highly structured nature of the cline width data, we built linear mixed models with nested random effects. We then tested whether the two key factors in the tension zone model - dispersal and selection against hybrids - predicted cline width, by specifying our dispersal estimates and mtDNA distance (our proxy for selection against hybrids) as fixed effects.

Our random effects model for the “all widths” data set accounted for the nested pattern of non-independence of the cline width estimates. Different cline width estimates are made 1) for the same hybrid zone transect, 2) for different transects but for the same hybridizing pair, and 3) for closely related taxa (phylogenetic non-independence). Thus, we include a random effect with levels corresponding to hybrid zone transect, which is nested within a random effect with levels corresponding to hybridizing pair. These random effects are further nested in a random effect with levels corresponding to clade at a broad taxonomic level (e.g. birds, frogs, insects; Fig. S1A).

Our random effects model assumes that the variance of cline width estimates is similar among levels of these random effects. However, speciation theory suggests that this variance should decrease as hybridizing pairs become more divergent (Barton 1983), which would result in heteroscedasticity of cline widths across mtDNA distances. We found no support for this prediction (*r*=−0.099, p=0.32; Fig. S8).

We additionally included a random effect for cline data type -- e.g. mtDNA markers, nuclear DNA markers, sex-linked genetic markers, morphological traits (Fig. S3C). As discussed above, cline widths can vary for both methodological and biological reasons, some of which are likely reflected by the type of character for which the cline is measured.

Finally, we expected that the relationship between mtDNA distance and selection against hybrids might vary across taxonomic groups, which may result in different relationships between mtDNA distance and cline width for different taxonomic groups. Accordingly, we attempted to estimate random slopes for mtDNA distance for each level (e.g. birds, frogs) of the taxonomic group. However, we dropped this term because it led to singularity issues. We instead investigated differences in relationships between mtDNA distance and cline width for each clade separately (see below).

Our full model included the nested random effect (transect within hybridizing pair within clade) and cline data type, and both fixed effects (dispersal, mtDNA distance) and their interaction. Prior to their inclusion, we took the natural log of both dispersal and mtDNA distance, and scaled them. We then evaluated model fit for the full model, and all possible simpler models, with respect to the fixed effects. Model fit was assessed using AICc scores. We report results from the best-fitting model and model-averaging done by AICc weights (Burnham and Anderson 2003). Additionally, we estimated the approximate *R^2^GLMM* for these models using the approach outlined in (Nakagawa and Schielzeth 2013).

### The “mean widths” approach

Then, we built a model in which we estimated a single cline width per hybrid zone. Variance in cline widths within a single hybrid zone can reflect differential introgression, stochastic variance, estimation error, and the use of non-diagnostic molecular markers to estimate clines. Our “all widths” approach does not explicitly account for these sources of variation, so including all widths for a given hybrid zone introduces unexplained variation. As we expect that each hybrid zone has a single geographic center where the transition between taxa is concentrated, we generated a point estimate for hybrid zone width by taking the geometric mean across all clines estimated for a given hybridizing pair. We analyzed this model in a linear model framework, including mtDNA distance, dispersal, and taxonomic group and all their pairwise interactions as predictor variables. We then evaluated model fit using AICc scores for the full model and all simpler models. We report results from the best-fitting model and model-averaging done by AICc weights.

Using the same data set, we conducted clade-specific analyses. Relationships between dispersal or mtDNA distance and hybrid zone width might be idiosyncratic across clades, and detecting such patterns is difficult in a clade-wide analysis. Such clade-specific patterns could be especially informative to understanding hybrid zone range boundary dynamics within particular taxa. To investigate evidence for clade-level patterns in our data, we fit linear models to the “mean widths” data from each clade with sufficient sampling to merit its own analysis: amphibians (n = 20), birds (n = 36), insects (n = 23), mammals (n = 27), and non-avian reptiles (n = 18).

### Sensitivity Analyses

We tested multiple variants of our model fitting to assess the robustness of our results. For our “all widths” data set, we considered four variants. First, our full data set includes both terrestrial and aquatic hybrid zone systems. As dispersal in aquatic systems can be directionally biased by abiotic factors (e.g. by gravity in river systems; or by ocean currents), these hybrid zone systems likely violate the tension zone model assumption that dispersal is isotropic. Terrestrial systems may be less prone to violating this assumption, so we performed an analysis limited to terrestrial systems.

Second, we assessed whether our approach to address phylogenetic non-independence might obscure the role of dispersal. In our “all widths” model, we included taxonomic group as a random effect to account for phylogenetic relatedness. However, taxonomic group also strongly predicts dispersal estimates (Fig. S2). Because some variance in cline width is attributed to taxonomic group instead of dispersal, we might underestimate the strength of the relationship between cline width and dispersal. Accordingly, we built a model in which we modeled taxonomic group, and its interaction with dispersal, as a fixed effect.

Third, we only considered clines based on mtDNA data. Given that our model used mtDNA distance as a proxy for selection against hybrids, we reasoned that mtDNA clines might best reflect this selection.

Fourth, we only considered those hybrid zones for which mtDNA distance was measured using the most often-used mitochondrial gene, *cytochrome b*. Considering just one gene allowed us to control for another unaccounted source of potential error: variance in substitution rates across mtDNA loci (Moritz et al. 1987; Eo and DeWoody 2010).

Using the “mean widths” data set, we then repeated model-fitting for a dataset containing only avian hybrid zones. For avian hybrid zones, we had alternative proxies for dispersal (HWI estimates) and selection against hybrids (phylogenetic distance) that were more standardized than the metrics used across all taxa in our analyses. As an alternative to dispersal estimates from the literature, we used the log of HWI (see *Methods: Dispersal estimates*). As an alternative to mtDNA distance, we used the time to MRCA in a large species-level phylogeny of birds (see *Methods: Divergence estimates*). Thus, birds -- which are also the most well-represented taxonomic group in our dataset -- offer an additional perspective on the robustness of our results.

All analyses were done in `R v3.3.3` using the statistical and graphing packages `ggplot2, cowplot, 1me4, r2glmm`, and `MuMIn` (Bates et al. 2007; Jaeger 2016; Wickham 2016; Wilke 2016; Barton and Barton 2018).

### Caveats of this work

The hybrid zones included in this study may be a biased sample of all likely hybridizing species pairs, which might limit the generality of these results. First, most of the hybridizing pairs included here are between morphologically well-defined taxa, while many less obvious hybrid zones likely exist between morphologically cryptic taxa. Thus, our sample is likely skewed towards more apparent hybrid zones, though it is unclear how this sampling bias would affect our results. Second, because measuring clines requires extensive population sampling through a hybrid zone, the hybrid zones included here must have had taxa with relatively high densities throughout the transition. Hybrid zones between taxa with lower densities may behave differently than those that can be adequately sampled for geographic cline analyses.

## Results

Our review of the hybrid zone literature identified 135 hybridizing taxon pairs for which we could find quantitative data on cline widths, dispersal estimates, and mtDNA divergence. Hybridizing pairs that we considered, but could not include due to incomplete data, are summarized in Table S3. Across hybridizing taxon pairs, we found data on a median of three clines per pair across an average of 2.1 cline types (Fig. S3B). The most common cline types measured were for nuclear and mtDNA markers (Fig. S3C). Hybridizing pairs occurred globally, though there was a strong bias towards studies from North America and Europe (Fig. S1B). Hybrid zones were fairly evenly split across vertebrate clades, though fish were underrepresented (Fig. S1A). Twenty-nine hybridizing pairs in our analysis were studied across more than one time point or geographical transect.

### “All widths” analyses

We investigated whether dispersal and mtDNA distance (a proxy for selection against hybrids) predict the spatial extent of hybrid zones. Our most inclusive analysis considered all estimated cline widths for each hybrid zone transect in a LMM framework (Fig. 3). Across all models, dispersal was supported as a fixed effect (model-averaged coefficient: 0.95 ± 0.16SE; relative importance (RI) = 1.0; Table 2; Fig. 3A) and mtDNA distance had limited support (coefficient: - 0.03 ± 0.9SE; RI = 0.25; Table 2; Fig. 3B). The interaction between dispersal and mtDNA distance had even weaker support (coefficient: 0.04 ± 0.1SE; RI = 0.13; Table 2). The best-fitting model only retained dispersal; dispersal explained *R^2^GLMM* = 0.16 of the variation in cline widths, while dispersal and the random effects together explained *R^2^GLMM* = 0.75 (Table 1).

**Figure 3.**
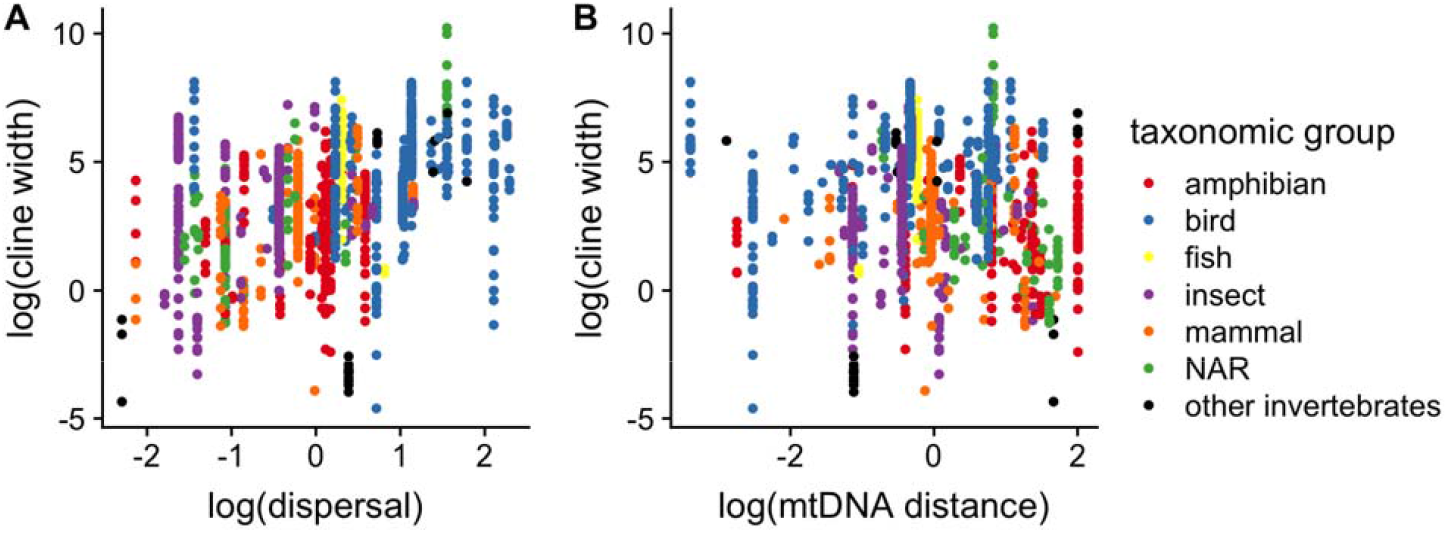
The relationship between cline width and (A) the log of dispersal and (b) the log of mtDNA distance for our “all widths” data set (*N* = 1488 clines across 131 hybrid zones). The best of our candidate models includes dispersal as the sole predictor (Table 1). NAR = non-avian reptiles.

**Table 1:** Model fitting for “all widths” data set (*N* = 1488 clines across 131 hybrid zones). Models predicting the log of cline width were fit using generalized linear mixed models. Predictors were the log of mtDNA distance and log of dispersal. Random effects were cline type (see Fig. S2C) and a series of nested random effects: transect within hybridizing pair within taxonomic group (see Fig. S1C). R^2^GLMM(m) shows the estimated proportion of variance explained by the fixed effects; R^2^GLMM(c) for both fixed and random effects. The best model is shown in bold.

| model | df | LnL | AICc | $\Delta\text{AICc}$ | $R^2\text{GLMM}(m)$ | $R^2\text{GLMM}(c)$ |
| --- | --- | --- | --- | --- | --- | --- |
| ~ log(mtDNA distance) × log(dispersal) | 9 | -2515.1 | 5048.4 | 3.6 | 0.154 | 0.749 |
| ~ log(mtDNA distance) + log(dispersal) | 8 | -2516.2 | 5048.4 | 3.6 | 0.159 | 0.753 |
| ~ log(mtDNA distance) | 7 | -2528.6 | 5071.3 | 26.5 | 0.007 | 0.759 |
| **~ log(dispersal)** | **7** | **-2515.4** | **5044.8** | **-** | **0.163** | **0.754** |
| ~ 1 | 6 | -2530.2 | 5072.4 | 24.2 | 0 | 0.761 |

**Table 2:**
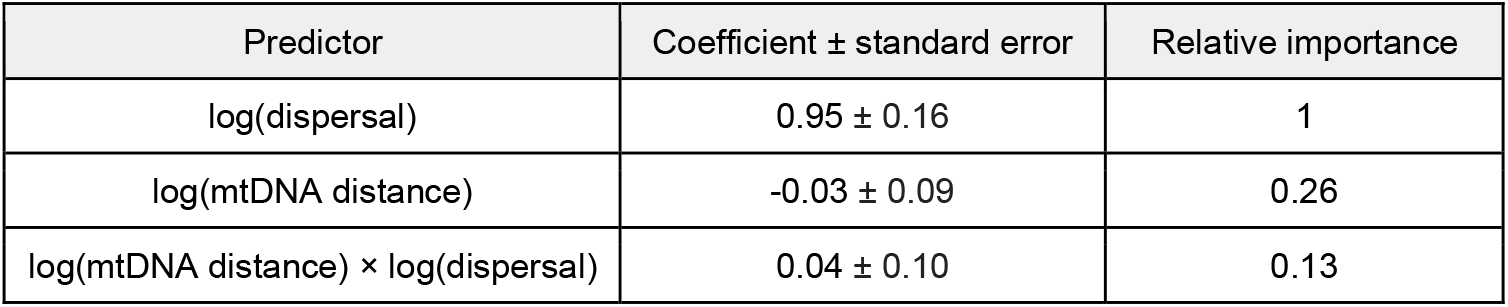
Model averaging results for the “all widths” models shown in Table 1, including coefficients and relative importance.

### “Mean widths” analyses

After finding the geometric mean cline width for each hybridizing pair, we fit the data set using a standard linear model framework (Fig. 4). Across all models, dispersal was supported as a predictor (model-averaged coefficient: 0.85 ± 0.41SE; relative importance (RI) = 0.99; Table 4; Fig. 4A) and mtDNA distance had very limited support (coefficient: −0.03 ± 0.11SE; RI = 0.49; Table 4; Fig. 4B). Taxonomic group and all interactions had even more limited support (Table 4). The best-fitting model only retained dispersal and had an adjusted *r^2^* = .31 (Table 3).

**Figure 4.**
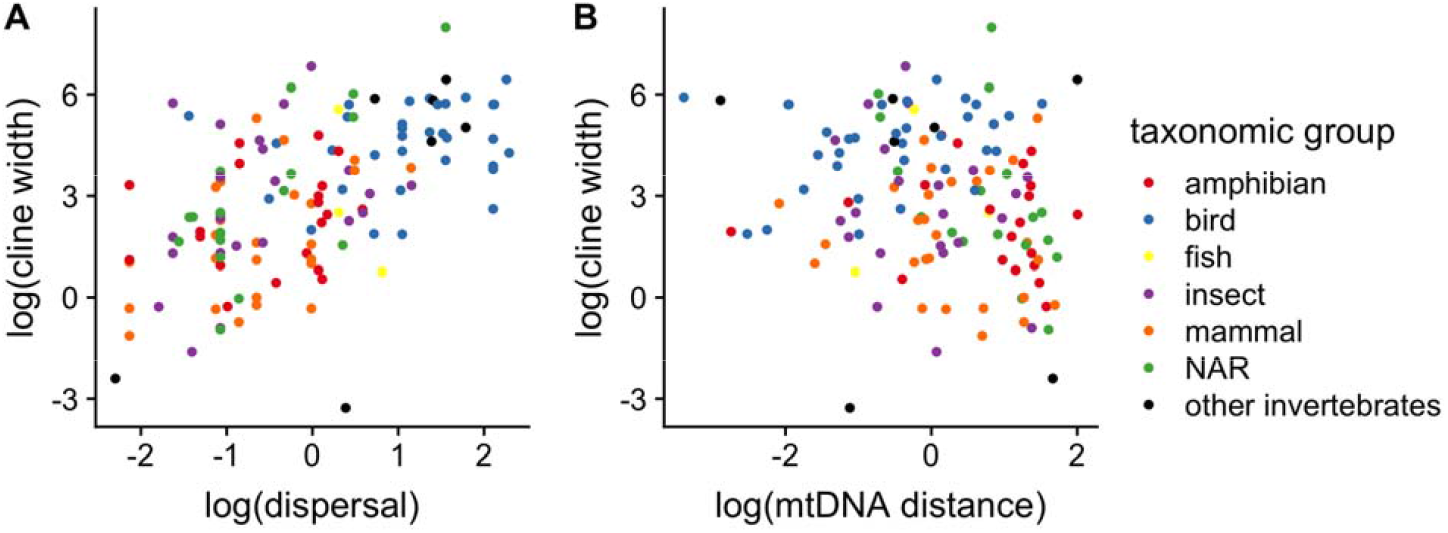
The relationship between mean cline width per hybrid zone and (A) the log of dispersal and (B) the log of mtDNA distance for our “mean widths” data set (*N* = 131 hybrid zone). All variables have been scaled. The best of our candidate models includes dispersal as the sole predictor (Table 3). The outlier with a highly negative residual is a hybrid zone between lineages of the coral species *Eunicea flexuosa*; removing this outlier did not result in qualitative changes in our results. NAR = non-avian reptiles.

**Table 3:** Model fitting for ‘mean widths’ data set, in which we calculated the geometric mean of cline width per hybrid zone (*N* = 131). We fit linear models that predicted the log of cline width, with the log of dispersal, log of mtDNA distance, taxonomic group, and two-way interactions as predictors. Relative likelihoods are calculated with respect to the model with the lowest AICc score. Shown are the five models with the highest relative likelihoods. The best model is shown in bold.

| model | AICc | adj. $r^2$ | rel. likelihood |
| --- | --- | --- | --- |
| **~ log(dispersal)** | **528** | **0.305** | **1** |
| ~ log(dispersal) + log(mtDNA distance) | 529.8 | 0.302 | 0.41 |
| ~ log(dispersal) × taxonomic group + log(mtDNA distance) | 530.2 | 0.378 | 0.33 |
| ~ log(dispersal) × taxonomic group + log(mtDNA distance) × log(dispersal) | 531.7 | 0.378 | 0.16 |
| ~ log(mtDNA distance) × log(dispersal) + taxonomic group | 533.1 | 0.329 | 0.08 |

**Table 4:** Model averaging results for the “mean widths” models shown in Table 3, including coefficients and relative importance. Coefficients not reported for predictors including taxonomic group because these are calculated for each one of the seven taxonomic groups.

| Predictor | Coefficient ± standard error | Relative importance |
| --- | --- | --- |
| ~ log(dispersal) | 0.84 ± 0.41 | 0.99 |
| ~ log(mtDNA distance) | -0.03 ± 0.11 | 0.46 |
| ~ taxonomic group | NA | 0.32 |
| ~ taxonomic group × log(dispersal) | NA | 0.24 |
| ~ log(mtDNA distance) × log(dispersal) | 0.02 ± 0.07 | 0.11 |
| ~ taxonomic group × log(mtDNA distance) | NA | 0.01 |

Our within-clade analyses recovered a positive relationship between dispersal and hybrid zone width, across all clades but amphibians, in which there was no relationship (Fig. 5, Table 5). However, the relationship between mtDNA distance and hybrid zone width varied among clades. Preferred models for birds and non-avian reptiles included mtDNA distance as a predictor of hybrid zone width. However, mtDNA distance and hybrid zone width scale positively in birds, while mtDNA distance showed a strongly negative relationship with hybrid zone width width for non-avian reptiles (NAR), as expected under the tension zone model (Fig. 5). Thus, the weak relationship between mtDNA distance and hybrid zone width from analyses across all animal taxa was partly explained by opposing patterns in different clades.

**Figure 5.**
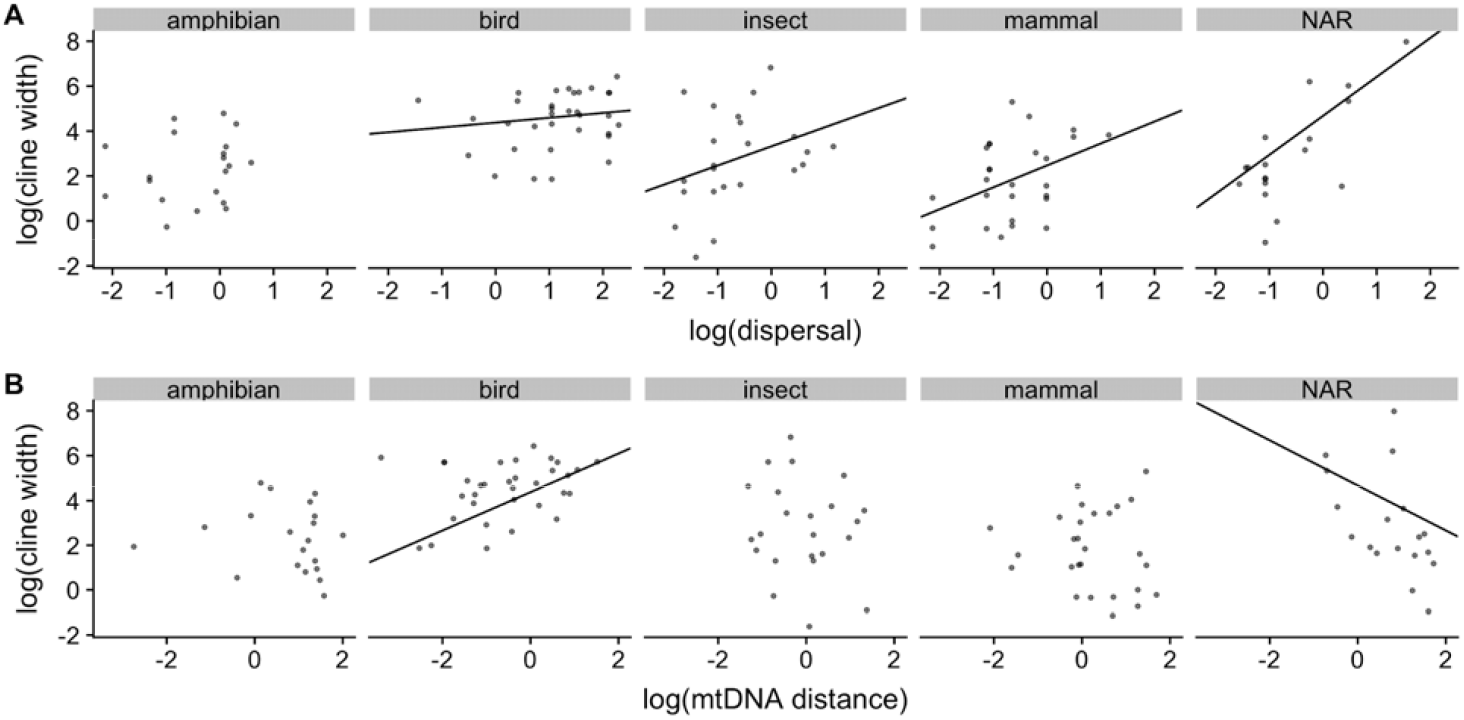
The relationship between cline width and (A) the log of dispersal and (B) the log of mtDNA distance, by taxonomic group. These data are shown for “mean widths” data set, in which we calculated the geometric mean of cline width per hybrid zone (*N* = 131). Shown are the five most well-represented taxonomic groups (NAR: non-avian reptiles). Also plotted are the linear fits for those variables that were included in the best-fitting model (Table 5). These data show that patterns vary substantially across taxonomic groups. While most groups show the predicted positive relationship between dispersal and cline width, only non-avian reptiles show the predicted negative relationship between mtDNA distance and cline width (Table 5).

**Table 5:** Results for the “mean widths” analysis, split by taxonomic group. We fit linear models that predicted the log of cline width by log of dispersal and of mtDNA distance within each taxonomic group shown. Shown are the details of the best-fitting model as determined by AICc scores. The predictors included in the best-fitting model are shown in the “Best model” column. Also shown are the estimated coefficients for all terms, and their relative importances, as determined by model averaging across the candidate model set.

| taxonomic group | *N* | Best model | $r^2$ | dispersal coefficient | mtDNA distance coefficient | Interaction coefficient | relative importance dispersal | relative importance mtDNA distance | relative importance interaction |
| --- | --- | --- | --- | --- | --- | --- | --- | --- | --- |
| amphibian | 20 | Intercept only | 0 | 0.08<br>(SE=0.27) | -0.04<br>(SE=0.18) | 0<br>(SE=0.06) | 0.26 | 0.23 | 0.01 |
| bird | 33 | ~ log(mtDNA distance) × log(dispersal) | 0.23 | 0.19<br>(SE=0.24) | 0.61<br>(SE=0.41) | -0.3<br>(SE=0.28) | 0.85 | 0.80 | 0.63 |
| insect | 23 | ~ log(dispersal) | 0.07 | 0.44<br>(SE=0.58) | -0.1<br>(SE=0.35) | 0<br>(SE=0.11) | 0.51 | 0.26 | 0.02 |
| mammal | 27 | ~ log(dispersal) | 0.16 | 0.74<br>(SE=0.52) | 0.11<br>(SE=0.34) | 0.43<br>(SE=0.76) | 0.88 | 0.45 | 0.30 |
| NAR | 18 | ~ log(mtDNA distance) × log(dispersal) | 0.61 | 1.79<br>(SE=0.51) | -0.75<br>(SE=0.61) | 0.01<br>(SE=0.2) | 0.99 | 0.75 | 0.09 |

### Sensitivity analyses

Using the “all widths” data set, we tested multiple variants of this basic model to assess the robustness of our results. These variants were: (1) inclusion of terrestrial systems only, (2) including taxonomic clade as a fixed rather than random effect, (3) inclusion of mtDNA clines only, and (4) inclusion of hybrid zones where mtDNA distance was measured using *cytochrome b* only. Across these four variants, we uncovered similar patterns to our primary “all widths” analysis. Model-averaging offered strong support for dispersal and much weaker support for mtDNA distance (Table S5, S7, S9, S11). The total portion of variance explained by the fixed effects in these models ranged from *R^2^GLMM*=0.069 - 0.325 (Table S4, S6, S8, S10).

Using the “mean widths” data set, we then fit models to avian hybridizing pairs only. As expected, our alternative proxies were correlated to the proxies used across all taxa (Fig. S4). Models using hand-wing index (HWI) as a proxy for dispersal support were qualitatively similar to analyses to analyses using dispersal estimates (Table S13, Fig. S6). Models with divergence time as a predictor of selection against hybrids provided less statistical support for an association between hybrid zone width and the two predictors, even after removing a single outlier (*Sula* boobies) (Table S15; Fig. S7).

## Discussion

Hybrid zones are a widespread phenomenon in animals. Our literature searches found 135 different hybrid zone-forming pairs that met the criteria for inclusion in our analyses (Table S3). The hybrid zones included in our study are a small subset of those found in nature. Most of these 135 hybrid zones are from temperate regions (Fig. S1B), suggesting that many hybrid zones wait to be discovered in the diverse tropics, where biotic factors like hybridization and competition are predicted to limit ranges more frequently than at high latitudes (Darwin 1859; Dobzhansky 1950; MacArthur 1972). Our meta-analysis underscores that hybrid zone formation is common, even among “good” species. The prevalence of hybrid zones in well-studied organisms indicates that hybrid zones are an important ecological phenomenon in the context of geographic range boundaries.

We found that random dispersal predicts hybrid zone widths across animals. Dispersal estimates alone explained >30% of the variation in estimated hybrid zone widths across our hybridizing taxon pairs. These hybrid zones occur across marine, freshwater, and terrestrial environments, and across a wide swath of diversity including invertebrate and vertebrate taxa. Moreover, we found evidence that dispersal predicts hybrid zone widths within four of the five taxonomic clades analyzed separately. Thus our results support dispersal as a major determinant of hybrid zone width on a global scale. This evidence, combined with the triangular shape of the relationship between dispersal and hybrid zone width (see below), suggests that directed dispersal at narrow ecological transitions infrequently results in narrow hybrid zones that map tightly to ecological gradients (or habitat transitions). Instead, it suggests that dispersal kernels do not regularly depart substantially from isotropic dispersal across hybrid zones. Consequently, the spatial scale of dispersal has consequences for the spatial scale of hybrid zones across animal diversity. However, as dispersal rate estimates explain only ~30% of the variation across hybrid zone widths via linear relationships on the log scale, we are left to explain the remaining fraction of the variation.

In the tension zone model, the shapes of range boundaries are stabilized when selection against hybrids counters dispersal across the hybrid zone. To examine support for this proposed balance between dispersal and selection, we tested whether mtDNA distance, as a proxy for selection against hybrids, explained variation in hybrid zone width. We found only limited support for this factor. In multiple analyses across the diversity of animals, we estimated a weakly negative relationship between mtDNA distance and hybrid zone width, with confidence intervals for coefficient estimates including zero. The tension zone model predicts a negative relationship, and while our results are consistent with this model, they provide weak support at most.

Why did we fail to recover strong evidence that increasing selection against hybrids -- here, measured as greater mtDNA distance -- leads to narrower hybrid zones? This could either be because mtDNA distance is a poor proxy for selection against hybrids, even while it is potentially a good proxy for smaller clades, e.g. non-avian reptiles (Fig. 5), or selection against hybrids does not strongly structure hybrid zones. If the former, then future studies may find that divergence in nuclear DNA or other, more direct measures of selection against hybrids are more predictive of hybrid zone width (Pereira et al. 2011). If the latter, we offer three explanations for the variation in our data. The first explanation is that selection against hybrids is not the primary force countering dispersal across hybrid zones. It is possible that other ecological forces - e.g. competition, pathogens (Case and Taper 2000; Case et al. 2005; Ricklefs 2010) - could play a strong role opposing dispersal across hybrid zones, with hybridization having limited ecological effects. Models suggest that in the absence of ecological gradients, these ecological processes would have to have greater inter-taxon than intra-taxon negative effects in order to stabilize the hybrid zone (Case et al. 2005; Goldberg and Lande 2007). Secondly, slight differences in fitness along ecological gradients, without habitat preferences, may be more important than selection against hybrids in stabilizing hybrid zone range boundaries (Kruuk et al. 1999).

Lastly, the efficacy of selection against hybrids in stabilizing hybrid zones depends on the rate of hybridization (or attempted hybridization). Tension zone models assume random mating, which ensures that hybrid zones will contain many offspring of mixed ancestry. Under these circumstances, both taxa will suffer negative demographic consequences due to the low fitness of many of their offspring. Assortative mating, however, can lessen these negative demographic effects: the fewer attempts at hybridization, the lesser the demographic consequences of selection against hybrids. If assortative mating is substantial in some fraction of hybrid zones, the relationship between hybrid zone width and selection against hybrids will break down, even if selection against hybrids is the primary force stabilizing the range boundary. Compared to random mating, assortative mating may allow a taxon to expand its range further into the other taxon’s range, widening the hybrid zone. Interestingly, we found positive relationships between mtDNA distance and cline width in clade-level analyses of birds, and to a lesser extent mammals (Table 5). This result raises the question of whether assortative mating, in addition to selection against hybrids, scales with mtDNA distance in some clades. If assortative mating scales with mtDNA distance, hybrid zone width may increase with mtDNA distance, instead of decreasing as in the tension zone model (Goldberg and Lande 2007).

Our analyses further allow us to identify which factors beyond dispersal are most important in structuring range boundaries set by hybrid zone dynamics. Specifically, there may be consistent explanations for positive versus negative residuals in our analyses. Positive residuals may correspond with hybrid zones undergoing neutral diffusion (Endler 1977), which may not ultimately be stable. Additionally, positive residuals may correspond with mosaic hybrid zones where habitats are substantially interdigitated, which may appear as clinal at large scale (Ross and Harrison 2002). In such cases, patches of habitat favorable to the locally rare taxon can stretch the width of the hybrid zone by supporting peripheral populations. Or, large, positive residuals from our analyses may correspond with hybrid zones exhibiting bounded hybrid superiority (Moore 1977), in which hybrids are equally or more fit than their parents. In this scenario, hybrid zone width should be determined by the spatial scale of the ecological transition over which hybrids are superior.

Hybrid zones with negative residuals with respect to dispersal may include instances where ecological gradients are steep, corresponding to strong selection, and/or where there is habitat choice. For example, the largest outlier in our “mean widths” analysis is from an extremely narrow hybrid zone between two lineages of the coral *Eunicea flexuosa*, which is illustrative of at least one, and perhaps both, of these scenarios (Prada and Hellberg 2014). *E. flexuosa* has broadcast dispersal, with population genetic dispersal rate estimates ranging from 2.9 to 55.52 km /√generation. However, the hybrid zone is less than 100 m in width. This zone corresponds with an extremely steep environmental gradient in depth and light availability, resulting in a steep selective gradient evident from reciprocal transplant experiments (Prada and Hellberg 2013) and age-class analyses (Prada and Hellberg 2014). It is additionally possible that despite their strong dispersal capacity, few *E. flexuosa* larvae settle in areas that poorly match their phenotypes (Figure 1 in (Prada and Hellberg 2014)), which could complement selection in narrowing the hybrid zone (Figure 1). Thus a combination of strong selection and habitat choice may yield the most potent departure toward narrowness from the pattern found for other hybrid zones.

While we report evidence for linear relationships between our two factors of interest and hybrid zone width, there are additional important patterns in these relationships. If we exclude the outlier *E. flexuosa* hybrid zone, the relationships between hybrid zone width and both dispersal and mtDNA distance are roughly triangular in shape, with an apparent lower bound (Figure 4). This shape underscores how rare narrow hybrid zones are in animals with strong dispersal (Prada and Hellberg 2014) or recent divergence, and is suggestive of limits on the narrowness of hybrid zones. Thus, even while hybrid zones are often narrow relative to taxon ranges, there may be limits to how narrow hybrid zones can be, imposed by dispersal and recency of divergence. If so, geography should limit where hybrid zones can form, particularly for dispersive and/or recently diverged taxa. For example, a hybrid zone between highly dispersive birds is unlikely to stabilize on a small island simply because space does not permit it. Rather, one taxon in the diverging pair is likely to outlast the other without hybrid zone formation. Because of these limitations, relatively few hybrid zones will stabilize in geographically restricted environments where space limits manifest. In evolutionary terms, our results reinforce the idea that there is a spatial scale of speciation (Kisel and Barraclough 2010) that may limit divergence processes. While extremely strong ecological gradients and habitat choice could lessen this restriction, the triangular shape of our data imply that these conditions either are rarely met, or that researchers have not interpreted existing instances as hybrid zones.

### Conclusions

We found that range limits set by hybridization are often narrow (Fig. 3, Fig. 4). To those who have studied hybrid zones for decades, the result might seem obvious. But for those of us who have not thought about hybrid zones as range boundaries, these results show how narrow geographic transitions between species can be, especially given that many of these hybrid zones form in the absence of an apparent environmental gradient (Brumfield et al. 2001; Singhal and Moritz 2012; McEntee et al. 2016). A previous summary found that the majority of hybrid zones were less than 50 times as wide as dispersal per generation (Barton and Hewitt 1985). Across these studies, we find that cline width for a hybrid zone is, at the median, 18.6□ greater than dispersal length. For example, in the iconic hybridizing pair of toads *Bombina bombina* and *B. variegata*, dispersal is estimated at 0.9 km / √generation. The hybrid zone between these two species is just 3.7 km wide (Szymura and Barton 1986, 1991). This pattern of narrow hybrid zones holds true even in species that disperse greater distances. In the hybridizing, migratory thrushes *Catharus ustulatus ustulatus* and *C. ustulatus swainsoni*, dispersal is estimated to be 150 km/√generation, and the hybrid zone is 71 km wide (Ruegg 2008). The narrowness of hybrid zones relative to range sizes suggests that many abrupt range boundaries might be explained in part by hybridization where other explanations like habitat transitions, strong gradients, or biogeographic boundaries have been invoked. For example, instances of elevational replacements, in which abrupt range boundaries are often assumed to result from local adaptation along steep environmental gradients, represent intriguing systems for exploration, as even these range boundaries may sometimes by stabilized by selection against hybrids, countering dispersal. As the tension model and our results suggest, narrow hybrid zones can form irrespective of whether or not these steep environmental gradient also result in a steep selective gradient (Fig. 4, (Goldberg and Lande 2007)).

Our results add more evidence for the importance of dispersal in determining range boundaries between species, both through species formation and the persistence of incipient or recent species. Dispersal is predicted to have contrasting but pivotal roles in species formation and species persistence. During species formation, too little dispersal makes it unlikely that population isolates will form (Mayr 1963). However, too much dispersal can retard divergence among isolated populations (Kisel and Barraclough 2010). Once species have formed, dispersal can facilitate sympatry by facilitating range expansion (Pigot and Tobias 2015; Weber and Strauss 2016; McEntee et al. 2018). However, if dispersal is too great, and species come into secondary contact before they evolve appreciable reproductive isolation (Levin et al. 1996; Weir and Price 2011), dispersal can also lead to the collapse of incipient species ranges. Our results show that greater dispersal increases the spatial extent of hybrid zones, which has implications for both species formation and coexistence. Additionally, we find that hybridizing species that disperse more also tend to be more genetically similar (as measured by mtDNA distance; Fig. S5) suggesting that hybrid zones are more likely to form earlier in the divergence history of high-dispersing taxa. However, we find no support for the premise that the age of hybridizing taxa structures the hybrid zone (Table 2 & 4).

This work focused on hybrid zones, in which hybridization assumes a defined spatial structure (Fig. 1). More broadly, other forms of hybridization can also play an important role in setting range limits. Generally, reproductive interference arising from hybridization -- whether via a hybrid zone or not -- can limit a species range (Case et al. 2005). However, determining when hybridization has limited species ranges can be difficult (Levin 2006). Because reproductive interference can lead to reproductive character displacement (Hoskin et al. 2005), taxa that are presently reproductively isolated could previously have interbred. Or, taxa can fail to establish because they go extinct via hybridization. In these cases, finding the “ghost of hybridization past” can be elusive (Levin 2006). That said, although challenging to do so, genomic data can reveal the occurrence of historical introgression, e.g., (Price et al. 2009; Martin et al. 2015). Analyzing these genomic data concordantly with the spatial scale of range limits might allow us to better understand how hybridization has constrained the evolution of range limits.

## Acknowledgements

Leo Shapiro and Catherine Sheard for sharing unpublished data. Rauri Bowie for making morphometric measurements on museum specimens at the Museum of Vertebrate Zoology. We thank Emily Lockwood and Hannah Hilbelink for assistance in making morphometric measurements. Florida Museum of Natural History for access and facilitation, especially Andrew Kratter and Thomas Webber. For thoughtful comments on earlier versions: Iris Holmes, Jimmy Peniston, Ricardo Pereira, Dan Rabosky. Funding from: NSF DEB-1519732 (to SS), NSF DBI-1458034 (to JGB), and the University of Florida.

## Data Accessibility

- Cline data and metadata from hybrid zones are in DataDryad Package: TBD
- Code used for data analysis & visualization are available from GitHub: TBD

